# Ethno-botanical Study of Medicinal Plants from Vidisha, Madhya Pradesh: Methanolic Extracts of Four Species and their Antimicrobial, Anti-inflammatory and Toxicity Properties

**DOI:** 10.1101/2023.05.08.539741

**Authors:** Urvashi Soni, Jitendra Malviya, Arvind Parmar, Diwakar Kr Singh

**Affiliations:** Department of Life Sciences and Biological Sciences, IES University, Bhopal; Department of Biotechnology, The Neotia University, Kolkotta (WB); Department of Pharmaceutical Sciences, SAGE University, Bhopal (M.P.)

**Keywords:** Rubia Cordifolia L, Caesalpinia crista L, Cassia alata L, and Plumbago zeylanica L, phytochemical, antioxidant, DPPH

## Abstract

The traditional usage of Plumbago zeylanica L, Cassia alata L, Caesalpinia crista L, and Rubia Cordifolia L in the villages of Vidisha district in Madhya Pradesh, India, has sparked research into the plants’ phytochemical makeup and a range of possible medicinal benefits. This research is the first to describe the phytochemical makeup and powerful anti-inflammatory and antioxidant properties of PZ stem. The PZ stem and CA fruit methanolic extracts had the greatest levels of phenolic, flavonoid, and flavonol compounds and had a robust antioxidant capacity for scavenging DPPH and hydrogen peroxide with regard to A. flavus, A. baumannii,, A. fumigatus, S. aureus, K. pneumonia, and R. oryzae, the antibacterial and antifungal activity of the chosen plant extracts were strongest for PZ, CA, and CA T. terrestris leaf.

## Introduction

Secondary metabolites such glycosides, saponins, flavonoids, steroids, tannins, alkaloids, and terpenes can be found in plants and may be used to combat pathogenic microbes [1]. These chemicals may have antibacterial or antifungal activity [2], and they are located in a variety of plant parts include seed roots, bark, leaves flowers & fruits. Many different diseases and infections can be treated with the same herb in traditional medicine [3]. Extracts from plants with a long past of outdated practice must be researched using contemporary techniques to examine their efficiency against human ailments in order to find novel medicines.

Though, rheumatoid arthritis is the primogenital known human ailment [4], finding a permanent solution for it has been difficult despite its prevalence across the globe. Although NSAIDs are widely used for pain and fever management, they are not effective against all forms of inflammation and have side effects include bleeding and ulceration [5]. However, secondary metabolites found in many plant species have been demonstrated to inhibit the growth of harmful microbes [1].

These metabolites contain glycosides, saponins, flavonoids, steroids, tannins, alkaloids, and terpenes.

Phytochemicals can be extracted from plant parts such roots, leaves, bark, flowers, fruits, and seeds to help combat bacterial and fungal infections [2]. Many different diseases and infections can be treated with the same herb in traditional medicine [3]. Extracts of plants that have a long history of traditional use must be researched using contemporary techniques to determine their efficiency against human ailments if we are to find novel medications.

This investigation centres on the Vidisha district of Madhya Pradesh, an area where Ayurveda has contributed numerous concepts for the medical application of plants. Traditional tribal tribes in the Vidisha area have long relied on the district’s ethnobotanically significant plants for their healthcare needs, and their comprehensive understanding of plants’ many facets makes the district an excellent place to study medicinal plants. A great number of tribal people, including those of the Bheel, the Sehariya, and the Mongiya, make their homes in this area, which is situated in the eastern part of the Malwa region. There are an estimated 1,458,875 people living in the Vidisha district as of the 2011 census, with 76.72 percent of those people calling rural areas home. The purpose of this ethnobotanical survey is to collect data on local medicinal plants that could be used in future studies.

In the foliage of host plants, you will often find tangles of fine, yellow-orange, heavily-branched stems from species like Rubia Cordifolia L (RC), Caesalpinia crista L (CC), Cassia alata L (CA), and Plumbago zeylanica L (PZ) [8]. Also found in this area is the annual noxious weed of crop fields known as Melilotus indicus (Fabaceae). Villagers in Madhya Pradesh’s Vidisha area rely on these plants as a primary line of defence against a wide range of infectious diseases.

Assays for lethality to brine prawns and cytotoxicity are utilised because they are well-established methods for detecting bioactivities in plant crude extracts. The use of plant extracts to test the toxicity of brine prawns before moving on to more involved antitumor and cytotoxicity experiments is an effective prescreening strategy. The cytotoxicity brine shrimp assay has been used in a number of studies to test plant extracts for antifungal, arthropod pest (including insect larvae), and molluscan activities, as well as for their anticancer and cytotoxic potential [11].

In this study, we wanted to see if certain plant extracts could indeed stop the spread of dangerous bacteria. The four medicinal plant species were examined for their phytochemical, antioxidant, antibacterial, cytotoxic, and anti-inflammatory properties.

Indian madder, or Rubia Cordifolia L (RC), is a medicinal herb with substantial scientific significance. Alizarin, a crimson pigment found in the plant’s roots, has been used as a natural dye for hundreds of years. In addition to its usage as a dye, RC has a long history of medical application in Ayurvedic practise for the treatment of conditions like skin inflammation, fever, and gastrointestinal distress.

Several research in recent years have looked on the possible health benefits of RC. Antioxidant, anti-inflammatory, and antibacterial activities were discovered in a root extract of the plant. Further, RC has demonstrated potential as a non-toxic treatment for liver disorders, cancer, and diabetes.

Traditional medicine based on RC has also been practised by a number of indigenous people groups and rural populations across India and the rest of Asia. The Bhil people of Rajasthan utilise the plant’s roots to cure snakebites, and the Kani people of Kerala use it to treat skin conditions. These historical use of RC have sparked contemporary studies that have improved our knowledge of the plant’s therapeutic qualities and newfound uses.

Several recent investigations have investigated the medicinal potential of Caesalpinia crista L in indigenous medicine. An extract of the plant’s leaves was found to have strong antioxidant and anti-inflammatory action in a 2019 study published in the Journal of Ethnopharmacology.

The plant’s hypoglycemic effects in animal models were also highlighted in a 2018 study that looked at the plant’s antidiabetic potential and was published in the same journal. A separate evaluation of the plant’s wound-healing qualities, published in the Journal of Pharmacy and Pharmacology in 2021, found that a topical ointment containing C. crista leaf extract significantly sped up the healing process and improved wound closure in animal models.

Recent studies have shown that Caesalpinia crista L. has considerable promise as a source of novel medications or natural products that can be included into indigenous medical practises.

The ringworm bush, or Cassia alata L, has long been revered as a medicinal plant among indigenous peoples. Recent studies have looked into its pharmacological and therapeutic potential.

Antimicrobial, antifungal, anti-inflammatory, antioxidant, and anti-diabetic effects have all been observed in studies with Cassia alata extracts. Infections, ringworm, scabies, and other skin problems have all been treated with it. It’s also been used to alleviate symptoms of diabetes and liver conditions.

Anthraquinones, flavonoids, and phenolic acids are just some of the bioactive chemicals isolated from Cassia alata, which are responsible for the plant’s therapeutic effects. The plant extracts have been studied extensively for their potential in alternative medicine and have been proven to have low toxicity and high safety profiles.

Recent studies on Cassia alata have revealed encouraging results, suggesting it may be a good place to find natural chemicals with therapeutic applications, especially for the treatment of skin infections and illnesses.

The medicinal properties of Plumbago zeylanica L. have been explored extensively. Recent studies have looked into its traditional benefits for treating inflammation, pain, diabetes, cancer, and microbiological infections.

Plumbago zeylanica L extracts, for example, showed strong anti-inflammatory and antioxidant effects in a study from 2020, suggesting they may be effective in the treatment of chronic inflammatory illnesses. Plumbago zeylanica L extracts were studied for their anti-diabetic potential in an animal model study published in 2021, and the results suggested that this plant could be a source for anti-diabetic medications.

Extracts of Plumbago zeylanica L. have been studied for their potential anti-cancer effects in several contexts. The potential of Plumbago zeylanica L extracts as an anti-cancer agent was shown by a 2021 study that demonstrated its ability to cause apoptosis (programmed cell death) in human lung cancer cells. Further validating its traditional use for treating microbial illnesses, a 2019 study demonstrated that Plumbago zeylanica L extracts displayed significant antibacterial activity against many types of bacteria and fungi.

Traditional usage in tribal medicine indicate that Plumbago zeylanica L. may be a useful source of medicinal ingredients for a wide range of ailments and disorders. To completely understand its modes of action and any adverse effects, however, more research is required.

## Methods

### Chemicals and plant material collection

The chemicals and reagents used in the study were purchased from Sigma-Aldrich, and included methanol, Folin-Ciocalteu reagent, terbinafine, streptomycin, bovine serum albumin (BSA), casein, perchloric acid, aspirin, Tris-HCl, and trypsin. Additionally, gallic acid, aluminum chloride, potassium acetate, sodium acetate, ascorbic acid, hydrogen peroxide, and phosphate buffer were used, all of which were of analytical grade.

In the Vidisha District of Madhya Pradesh, India, samples of the plant species Rubia Cordifolia L (RC), Plumbago zeylanica L (PZ)(stems), Caesalpinia crista L (CC), and Cassia alata (CA) and leaf extracts (CA) were gathered from close by villages. Dr. Ajay Bharadwaj, Professor at the Institute of Biotechnology at IEHE in Bhopal, identified the plants. The plant species were chosen based on information from locals and traditional herbalists that the extracts are mostly used to cure various infectious disorders.

**Table 1.** list the four-plant species’ scientific, English, and regional names as well as the sections of the plants customarily extracted, their ethnomedicinal use, administration methods, and extract yields.

| S.No. | Botanical name, family | Parts used | Ethno medicinal use | Route of administration | Yield (%) |
| --- | --- | --- | --- | --- | --- |
| 1 | *Rubia Cordifolia* L (RC) | Leaves | Gastrointestinal disorders, skin diseases, fever, diabetes, and rheumatism.<br>Digestive, an appetizer, and a blood purifier. | Topical, Oral | 31 |
| 2 | *Caesalpinia crista* L (CC) | Stem | As fever, inflammation, pain, and skin diseases. As a tonic and astringent, as well as for the treatment of respiratory tract infections, bronchitis, and asthma. Traditionally used for its antidiabetic and hepatoprotective properties. Decrease blood glucose levels and improve lipid profiles in diabetic animal models. Reducing liver damage and increasing antioxidant levels. | Oral, Topical | 17 |
| 3 | *Cassia alata* L (CA) | Leaves | used to treat skin infections, wounds, fungal infections, and snake bites. Eczema and psoriasis, and to treat ringworm and other fungal infections. Snake bites, and the bark is used to treat dysentery and diarrhea. Treat fever, cough, and respiratory infections, as well as gastrointestinal disorders, such as constipation and indigestion. It is believed to have antimicrobial, anti-inflammatory, and analgesic properties. | Oral, Topical | 29 |
| 4 | Plumbago zeylanica L (PZ) | Leaves | Digestive disorders, respiratory problems, skin diseases, and rheumatism. Snake bites and scorpion stings, remedy for toothache and oral diseases. To treat mouth sores. | Oral, Topical | 37 |

### The process of making plant extracts

The leaves, stems, and fruits of the selected plant species were washed thoroughly with distilled water and left to dry for three days at room temperature in the shade. The dried plant material was then equally ground using an electric grinder. 250 g of the resulting powder were soaked in one litre of 100% methanol for four days [12]. The methanol extract was fully dried in a rotary evaporator at room temperature (30 °C) after filtering using Whatman No. 1 filter paper. After drying at ambient temperature, the thick extract was then refrigerated at 4 °C for subsequent use. The percentage yield (w/w) of the dried plant extract was calculated by dividing the weight of the dried extract by the weight of the dried plant material and multiplying the result by 100. The use of analytical-quality chemicals and reagents throughout the method must be emphasised.

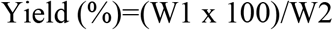

Where W2 is the weight of the soaking plant powder and W1 is the dry weight of the extract after the solvent has evaporated.

#### Preliminary phytochemical analysis

To determine the presence of certain phytochemical components such saponins, terpenoids, anthraquinones, phlobatannins, flavonoids, and phenolic compounds, the chosen plant extracts were submitted to phytochemical screening [12–14].

#### Thin-layer chromatography

Capillary tubes were used to apply the 10 L of methanolic plant extracts on TLC plates that had already been coated before being given time to RCdry. The TLC plates were developed in a chamber using chloroform:methanol (5:1) as the mobile phase, and they were afterwards inspected under UV light. Caffeic acid, quercetin, rutin, trans-cinnamic acid, and salicylic acid were considered standards.

The mobility of the samples was expressed by the retention factor (Rf), which was calculated using the method below:

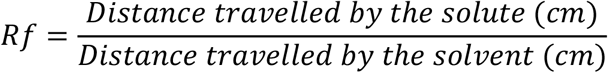

### Quantitative analysis of phytochemicals

The Folin-Ciocalteu colorimetric technique was modified to determine the total phenolic content. An aliquot of the plant extract was quickly combined with the Folin-Ciocalteu phenol reagent and let to stand for 5 minutes at room temperature. The mixture was then let 90 minutes to stand at room temperature after the addition of 6% sodium carbonate. Using a spectrophotometer, the mixture’s absorbance was determined at 725 nm. The same process was used to create a calibration curve for gallic acid in the concentration range of 20–80 g/mL. Results were given as mg of gallic acid equivalent (GAE) for every gramme of extract.

The aluminium chloride colorimetric technique was modified to determine the total flavonoid concentration. Briefly, 10% aluminium chloride, 1 M potassium acetate, 80% methanol, and distilled water were combined in separate test tubes with plant extract and standard. After 30 minutes of room temperature incubation, the mixture was measured at 415 nm for absorbance. The amount of flavonoid per gramme of extract was represented as mg quercetin equivalent (QE). For quercetin concentrations between 20 and 80 g/mL, a calibration curve was created.

With slight adjustments, the aluminium chloride colorimetric technique was used to calculate the total flavonol concentration. In a nutshell, 2% aluminium chloride and 5% sodium acetate were combined with the extract and standard in individual test tubes. After centrifuging the mixture to produce a clear solution, the absorbance was measured at 440 nm. Per gramme of extract, the findings were represented as mg quercetin equivalent (QE). For quercetin concentrations between 20 and 80 g/mL, a calibration curve was created. Three duplicates of each plant extract were created.

#### Activity to scavenge DPPH radicals

The 2,2’-diphenyl-1-picrylhydrazyl (DPPH) test was used to gauge the plant extracts’ in vitro free radical scavenging capacity [20, 21]. Three millilitres of methanol, one millilitre of extract, and one millilitre of DPPH (0.3 mM) in methanol made up the reaction mixture. We tested the scavenging activity of the materials at different doses (25-200 g/mL). The reaction mixture was shook vigorously after being incubated for 10 minutes in the dark at room temperature. At 517 nm, absorbance was measured. In the same way as the samples, the control was made, but without the plant extracts. Rutin [23] and ascorbic acid [22] were employed as positive controls. The fraction of DPPH radical scavenged was computed based on the following equation: “

Effect of scavenging (%) = [(control absorbance-sample absorbance)/control absorbance] X 100 The approach used to evaluate the hydrogen peroxide-scavenging capacity of plant extracts was adopted from Ruch et al. (1989) [24]. In phosphate buffer (50 mM, pH 7.4), hydrogen peroxide (2 mM) was produced as a solution. Using 50 mM phosphate buffer (pH 7.4), aliquots of plant extracts (0.1 mL) were placed into vials and their quantities increased to 0.4 mL. Plant extracts (25–200 g powder/mL) were produced in distilled water. After 0.6 mL of hydrogen peroxide solution was added, the tubes were vortexed, and after 10 minutes, the absorbance was measured at 230 nm in comparison to a control solution made up of phosphate buffer without hydrogen peroxide. Rutin and ascorbic acid were utilised as positive controls. The following equation was used to determine the extract’s capacity to scavenge hydrogen peroxide:

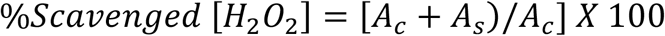

Staphylococcus aureus (ATCC 6538), Pseudomonas aeruginosa (ATCC 7221), Klebsiella pneumonia (ATCC 6059), Aspergillus flavus (ATCC 22456), Aspergillus fumigatus (ATCC 26934), and Rhizopus oryzeae (ATCC 11886) were the bacterial and fungal strains used in this study. The Department of Life Sciences and Biological Sciences at IES University in Bhopal as well as the ATCC provided these strains.

The fungal strains were sub-cultured on potato dextrose agar (PDA) Hi-media, Mumbai for 7 days at 25 °C, while the bacterial strains were sub-cultured in nutritional broth (Sigma) and incubated at 37 °C for 18 hours.

The antibacterial and antifungal assays used terbinafine (1 mg/mL) and streptomycin (30 g/mL) as positive controls. For the assessments of the antibacterial and antifungal effects, DMSO was employed as a negative control.

The absorbance of the control (AC) and sample (AS) was determined using the techniques described in the previous paragraphs in order to evaluate the antibacterial and antifungal properties of the plant extracts. test for antibacterial effectiveness

The agar well diffusion technique was used to assess the antibacterial activity of methanol extracts from the selected plant species [26]. 75 mL of seeded MH agar was put onto the Petri plates, and the agar was let to harden. A sterile cotton swab was then used to uniformly distribute the bacterial inoculum throughout the whole agar surface. A well was made and filled with 100 l of each crude extract using a sterile cork borer (6 mm). The Petri plates were incubated at 37 °C overnight after being maintained at room temperature for an hour. The control plates’ well was filled with solvent. The plates were checked for zones of inhibition after 24 hours, and the findings were compared to those of the positive control, streptomycin (30 g/mL), to see how they fared.

Calculation of the Minimum Bactericidal Concentration (MBC) and the Minimum Inhibitory Concentration (CCC)

The microdilution technique [27] was used with nutritional broth to determine the minimum inhibitory concentration (MIC) and minimum bactericidal concentration (MBC) of the methanolic extracts from the selected plant species. The plant extracts were dissolved in 10% DMSO, and culture broth was used to make two-fold dilutions. Each test sample and growth control was injected with 10 l of a bacterial solution containing 5 106 CFU/mL. The growth control comprised broth and DMSO without plant extract or antibiotic agent. Each sample also received a resazurin solution (270 mg of resazurin tablet diluted in 40 mL of sterile water). The samples were then incubated at 37 °C for 24 hours. The absorbance at 500 nm was measured in order to identify bacterial growth. Bacterial growth was seen as a shift in colour from purple to pink or colorlessness. The MIC was defined as the lowest concentration of plant extract that changed colour or the greatest dilution that totally prevented bacterial growth. The test dilutions were further sub-cultured on new solid medium and incubated for 18 hours to ascertain the MBC values. MBC was defined as the maximum dilution of plant extract at the lowest concentration necessary to eradicate all bacteria. The CCC and MBC values for each dilution were evaluated in triplicate for each bacterial strain.

test for antifungal effectiveness The agar tube dilution technique was used to assess the antifungal activity of the chosen plant species’ methanol extracts [28]. The unprocessed plant extract was dissolved in DMSO to create the samples. 6.5 g of potato dextrose agar were dissolved in 100 mL of distilled water (pH 5.6) to create culture medium. Test tubes with cotton plugs or screw-capped caps were filled with potato dextrose agar (10 mL), and the tubes were autoclaved at 121 °C for 21 minutes. The potato dextrose agar was filled with 67 L of extract from the stock solutions and the tubes were allowed to cool at 50 °C. The media-filled tubes were then allowed to harden at room temperature while tilted. A 4-mm-diameter piece of inoculum collected from a 7-day-old fungal culture was used to inoculate the tubes containing solidified medium and plant extract. Parallel controls were carried out using the appropriate solvent in place of the plant extract. The cultures were checked twice a week while the test tubes were cultured for 7 days at 28 °C. The fungus in the slant was measured for linear length (mm), and growth inhibition was computed using the negative control. The studies were carried out in triplicate, and the percentage of fungal growth inhibition at each chemical concentration was calculated.

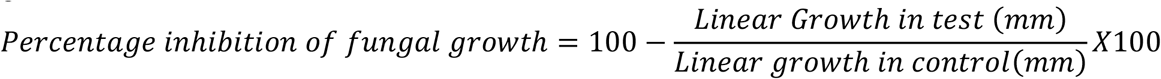

Using the nauplii of brine shrimp, the cytotoxic activity of the methanolic plant extracts was assessed [29]. The chosen plant extracts were dissolved in DMSO to produce 10, 100, and 1000 mg/mL final concentrations. Each concentration was divided into two millilitres, transferred to a graduated vial, evaporated to dryness for 48 hours, and then reconstituted with DMSO. The adult nauplii of brine shrimp were collected after the eggs had hatched for 48 hours in a plastic container with aeration. Each vial received ten nauplii, and 2 mL of saline solution was added to form the final volume. The vials were incubated for 24 hours at 25 °C, and the number of nauplii that survived were counted using a 3 magnification. Potassium dichromate served as the reference standard, while DMSO and saline solution served as the negative controls. Through the use of linear regression analysis, the fatal dosage was determined [30]. The Abbott formula was used to determine the fatality percentage.

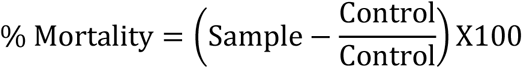

### Anti-inflammatory activity

#### Inhibition of protein denaturation method

We used the approach described by Tizushima et al. [31], with a few minor adjustments, in order to assess whether or not a substance inhibits the denaturation of proteins. In the reaction mixture, 1% BSA was present as an aqueous solution, and various quantities of the test extract were also present. To achieve the desired pH level in the reaction mixture, 1N HCl was used. After being heated at 37 degrees Celsius for twenty minutes, the samples were then heated at 57 degrees Celsius for twenty minutes before being allowed to cool. At a wavelength of 660 nm, the turbidity of the samples was evaluated. The experiment was carried out three times for accuracy. The following is the calculation that was used to get the percentage of inhibition of protein denaturation:% Inhibition = (AC - AS) / AC x 100

Where, AC and AS are the absorbance (at 600 nm) of the control and sample, respectively.

#### Human red blood cell (HRBC) membrane stabilization test

Fresh human blood of a volume of 10 millilitres was drawn into heparinized centrifuge tubes and then spun at a speed of 3000 revolutions per minute for ten minutes. This was done so that the membrane-stabilizing activity of the methanolic plant extracts could be evaluated. After measuring the volume of the blood, which was then reconstituted as a 10% v/v suspension with normal saline, the red blood cells were washed three times with normal saline solution [32]. It was decided to make a reaction combination of two millilitres by combining one millilitre of the plant extract in methanol with one millilitre of the 10% red blood cell suspension. In addition, a saline-based “control” solution that did not include any plant extract was made. In this experiment, the “positive control,” or “standard drug,” was aspirin. After incubating the samples for 30 minutes at 56 degrees Celsius and centrifuging them at 2500 revolutions per minute for five minutes, the absorbance of the supernatant was measured at 560 nanometers. The experiment was carried out three times to ensure accuracy.

The formula that was used to compute the percentage of membrane stabilisation activity was presented in the part titled “Inhibition of protein denaturation method” [33], whereas the formula that was used to calculate the percentage of protection was presented as follows:

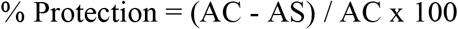

Where AC and AS represent the absorbance (at 560 nm) of the control and sample, respectively.

#### Proteinase inhibitory assay

The Proteinase inhibitory test was carried out using a methodology that had been altered from that described by Oyedepo and Femurewa [34]. In the reaction mixture that was 2 millilitres in volume, there was 0.06 mg of trypsin, 1 millilitre of Tris-HCl buffer with a pH of 7.4, and 1 millilitre of test plant extract sample with varying quantities. After heating the reaction mixture to 37 degrees Celsius for five minutes, one millilitre of casein with a weight-to-volume ratio of 0.8% was added. A total of twenty more minutes were spent incubating the combination. To halt the process, two millilitres of perchloric acid with a concentration of seventy percent was added. After centrifuging the hazy solution, the absorbance of the supernatant was measured at 210 nm against a Tris-HCl buffer serving as a blank. The experiment was carried out three times for accuracy.

Following the completion of the statistical analysis, the findings were reported using the mean along with the standard error of the mean (SEM). A one-way analysis of variance (ANOVA) was performed using Statistic version 8.1 on the data that was produced from quantitative testing for phytochemicals and antifungal activity. The Least Significant Difference (LSD) test was used to determine whether or not there were significant differences between groups when P was less than 0.05 [35]. The IC50 values were determined by the use of linear regression analysis. The R programme was used to do an analysis of linear correlations using regression (3.2.2.).

## Results

A qualitative study of phytochemicals, TLC, and measurement of the total contents of phenolic, flavonoid, and flavanol were carried out. The investigation revealed that the extracts of all of the plants under consideration included phenolics, flavonoids, and terpenoids. Tannins were discovered in CC, RC, and PZ, although phlobatannins were only discovered in PZ and CAF. Tannins were discovered in all three locations. CAL was found to have the greatest levels of anthraquinone, whereas RC had the lowest levels (Table 2). Using a technique called thin-layer chromatography, we were able to validate the presence of a variety of bioactive chemicals in each plant extract (Table 3). The extracts of RC’s leaves included a total of six spots, but the extracts of PZ’s stems had a total of nine spots. The TLC analysis of CC leaf extracts showed six spots, while the TLC analysis of CA leaf extracts also produced six spots. According to the results of the quantitative study, PZ had the greatest levels of phenolic, flavonoid, and flavanol content (497, 385, and 139 mg/g, respectively), followed by CA (426, 371, and 138 mg/g). The lowest levels of phenolic and flavonoid compounds were found in CC and RC, respectively, as shown in Table 4.

**Table 2.**
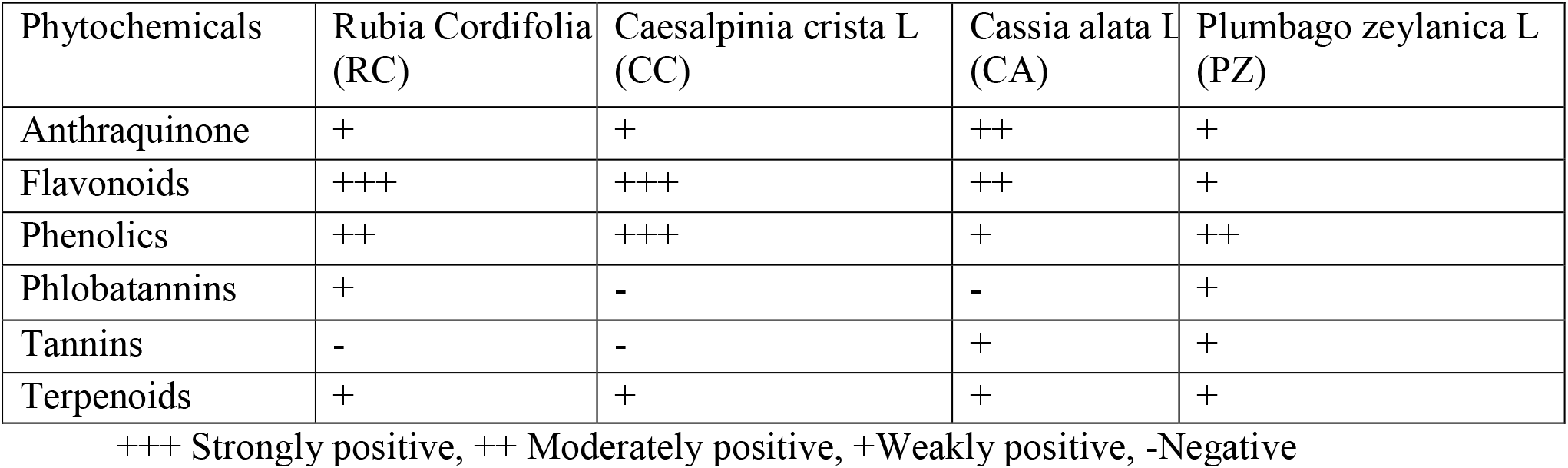
Antioxidant activities.

| Phytochemicals | Rubia Cordifolia (RC) | Caesalpinia crista L (CC) | Cassia alata L (CA) | Plumbago zeylanica L (PZ) |
| --- | --- | --- | --- | --- |
| Anthraquinone | + | + | ++ | + |
| Flavonoids | +++ | +++ | ++ | + |
| Phenolics | ++ | +++ | + | ++ |
| Phlobatannins | + | - | - | + |
| Tannins | - | - | + | + |
| Terpenoids | + | + | + | + |

+++ Strongly positive, ++ Moderately positive, +Weakly positive, -Negative

**Table 3.** The potential of plant extract.

| Potential Standard | (R<sub>f</sub>) Value |
| --- | --- |
| Caffeic acid | 0.85 |
| Quercitin | 0.38 |
| Rutin | 0.97 |
| *trans*-cinnamic acid | 0.74 |
| Salicylic acid | 0.6 |
| Plant extracts | R<sub>f</sub> value |
| Rubia Cordifolia L (RC), and | 0.23, 0.26, 0.31, 0.45, 0.56, 0.83 |
| Caesalpinia crista L (CC), | 0.16, 0.2, 0.25, 0.35, 0.45, 0.61, |
| Cassia alata L (CA) | 0.75, 0.83, 0.85 |
| Plumbago zeylanica L (PZ) | 0.20, 0.25, 0.34, 0.37, 0.56, 0.83 |

**Table 4.**
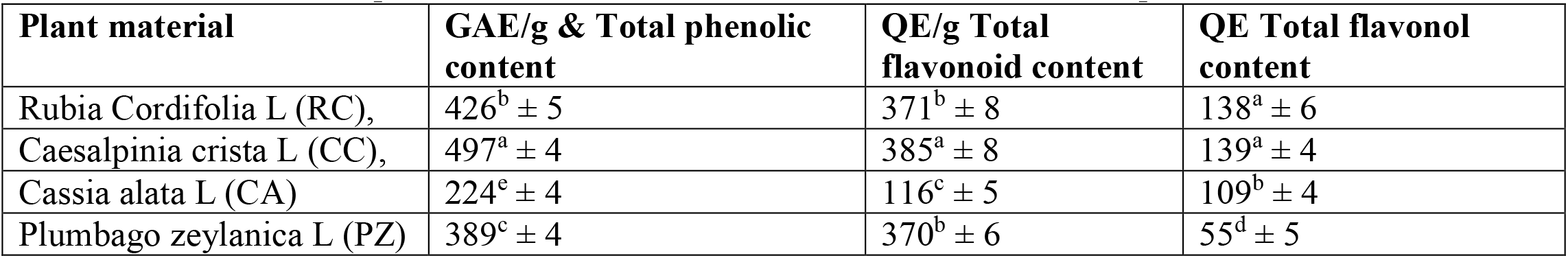
Total phenolics, flavonoid and flavonol content in the dried plant extracts

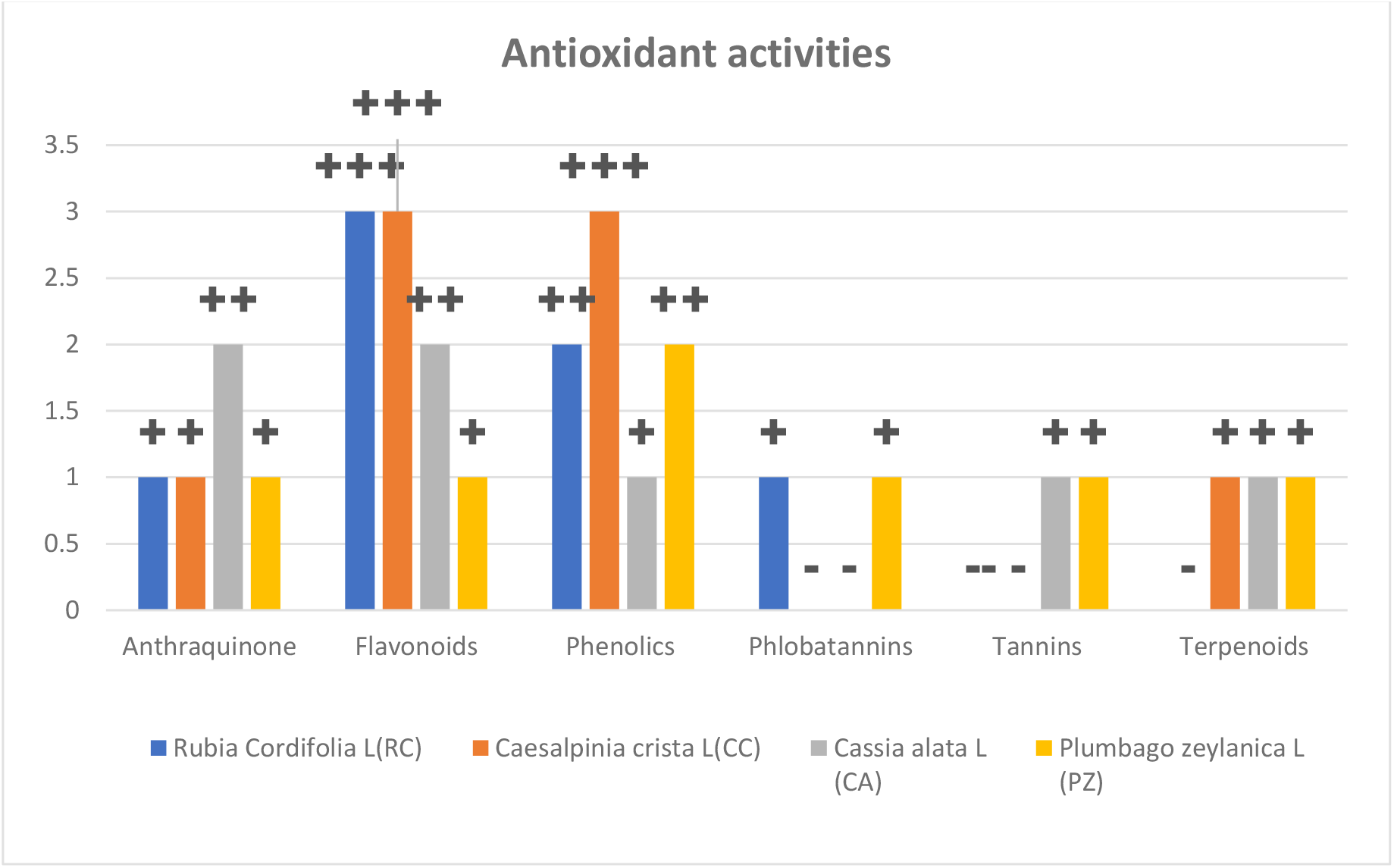

The DPPH test is a method that may be used to assess whether or not plant extracts or compounds have the capacity to serve as free radical scavengers and have antioxidant potential. PZ and CA extracts were tested for their ability to neutralise DPPH radicals, and the results showed that their respective IC50 values were 20 1 and 92 2 g/mL. On the other hand, their capacity to scavenge free radicals was significantly inferior to that of ascorbic acid and rutin (5 0.4, 18 0.5 g/mL). In terms of their ability to remove DPPH from the body, the chosen plant extracts were rated as follows: PZ > CA > CA > RC > CC. PZ and CAF extracts showed substantial hydrogen peroxide radical scavenging activity (IC50 values of 18 0.7 and 26 2 g/mL, respectively). Additionally, the IC50 values of the chosen plant extracts for H2O2 scavenging activity were obtained. Using the agar well diffusion technique, the antibacterial activity of the plant extracts was examined to see how they fared against a variety of different microbes. P. aeruginosa, S. aureus, K. pneumoneae, and A. baumannii were all significantly inhibited by the PZ extract, which also had the lowest MIC and MBC values. An extensive investigation into the relationship between the total phenolics, DPPH, and H2O2 activities of the five plant extracts resulted in the discovery of a high correlation coefficient (R2) of 0.93.

Antifungal activityPresented in figure 3 are the findings of the tests conducted to evaluate the antifungal capability of the various plant extracts. The mycelial growth of A. flavus was most effectively inhibited by methanolic extracts of PZ, which achieved an 88% inhibition rate, followed by CA and CA, which achieved 83% and 79% inhibition rates, respectively. In terms of the antifungal potential of plant extracts against A. flavus, the order of PZ > CA > CA > CC > RC was shown to be most effective. PZ extract showed the greatest decrease (90%) in the mycelial growth of A. fumigatus, followed by CA extract (87% reduction) and CA extract (85% reduction), respectively. PZ was at the top of the list when it came to the antifungal potential of plant extracts against A. fumigatus, followed by CA, CC, and RC in that order. PZ was once again the most effective treatment for inhibiting the mycelial development of R. oryzae (94%); the next best treatments were CA and CA (93% and 90%, respectively).

**Figure 1.**
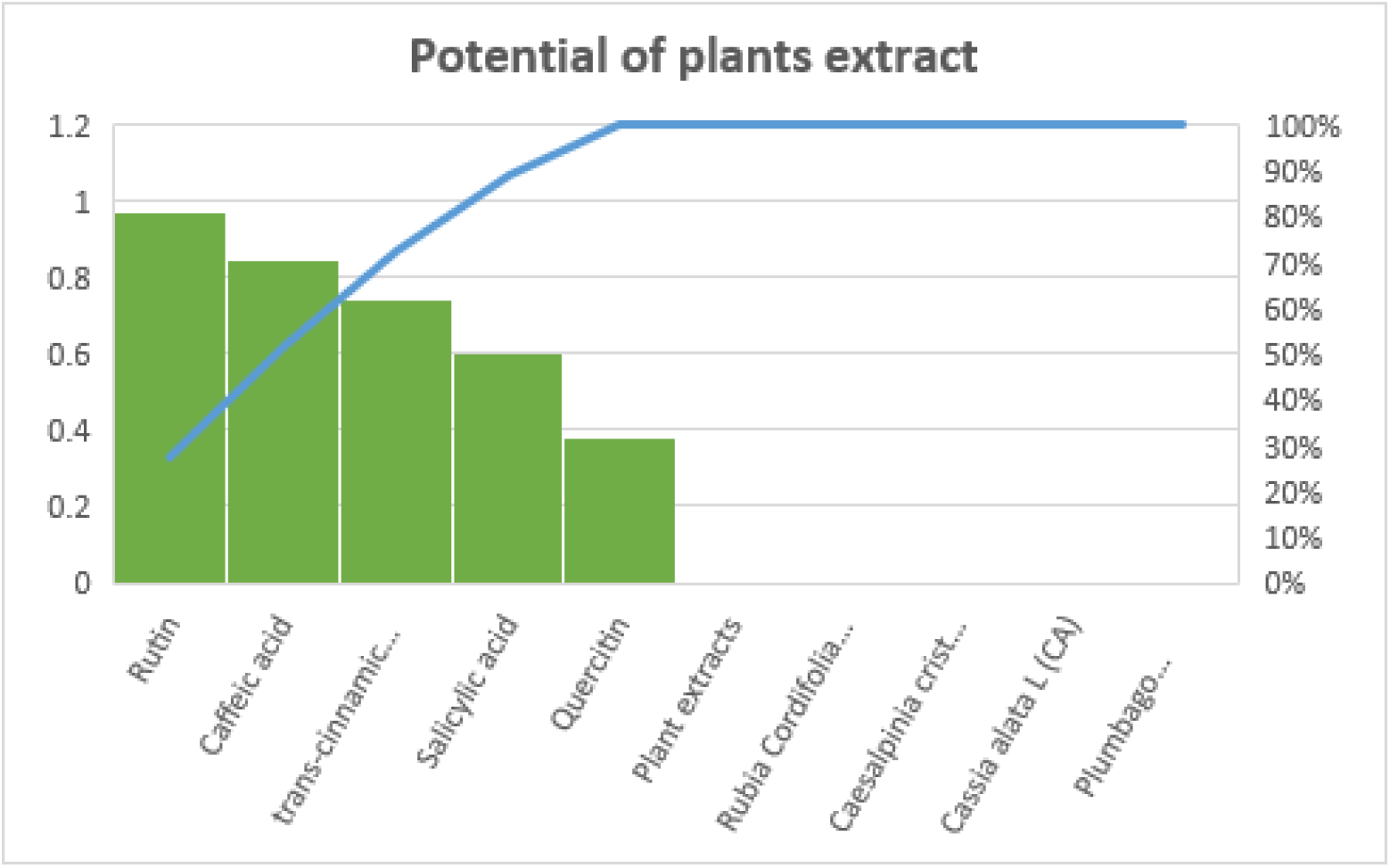
Plant Extract potential

**Figure 2.**
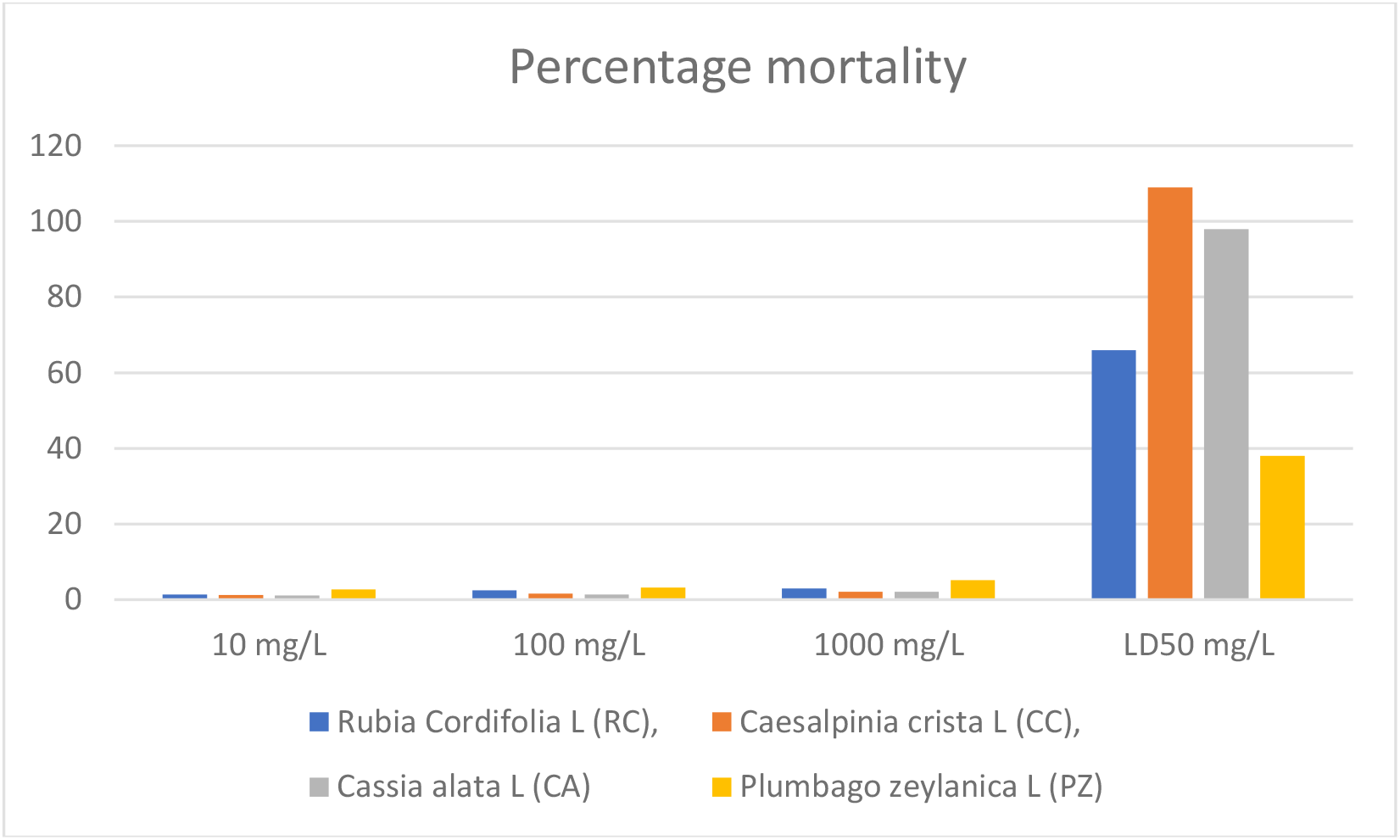
Mortality Percentage

**Figure 3.**
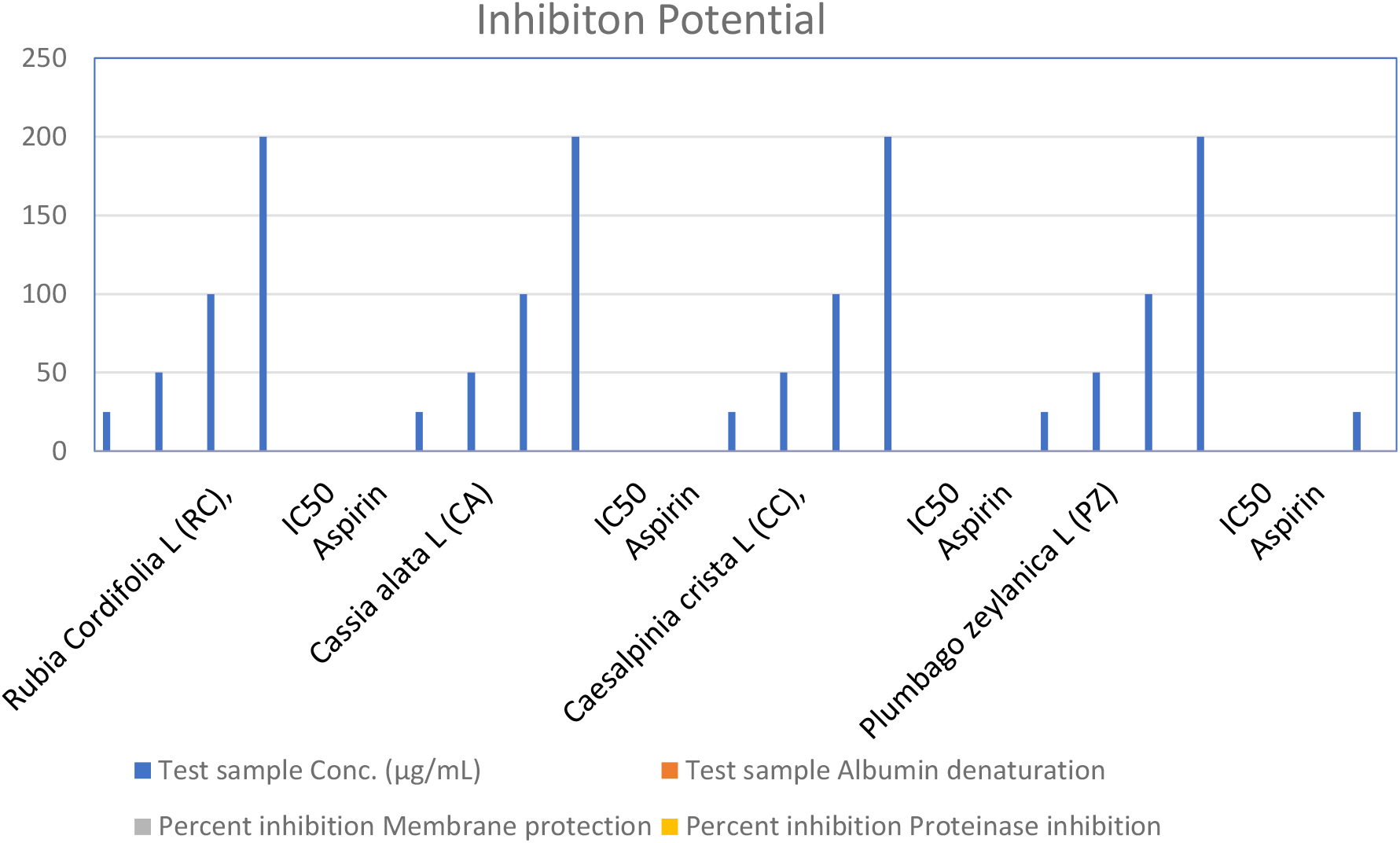
Linear correlation of the total phenolic content versus the anti-inflammatory and anti-proteinase activity of selected plant extracts

Antimicrobial activity have been shown to have a correlation with total phenolic content. There were found to be positive connections between the amount of phenolic compounds present in the chosen methanolic plant extracts and the amount of bacteria and fungus that were inhibited (Fig. 4a, b). It was discovered that the correlation coefficients (R2) for activity against A. baumannii, K. pneumoneae, P. aeruginosa, and S. aureus were respectively 0.55, 0.66, 0.87, and 0.46. The R2 values for total phenolic content and antifungal potential were 0.92, 0.70, and 0.53 against A. flavus, A. fumigatus, and R. oryzae, respectively.

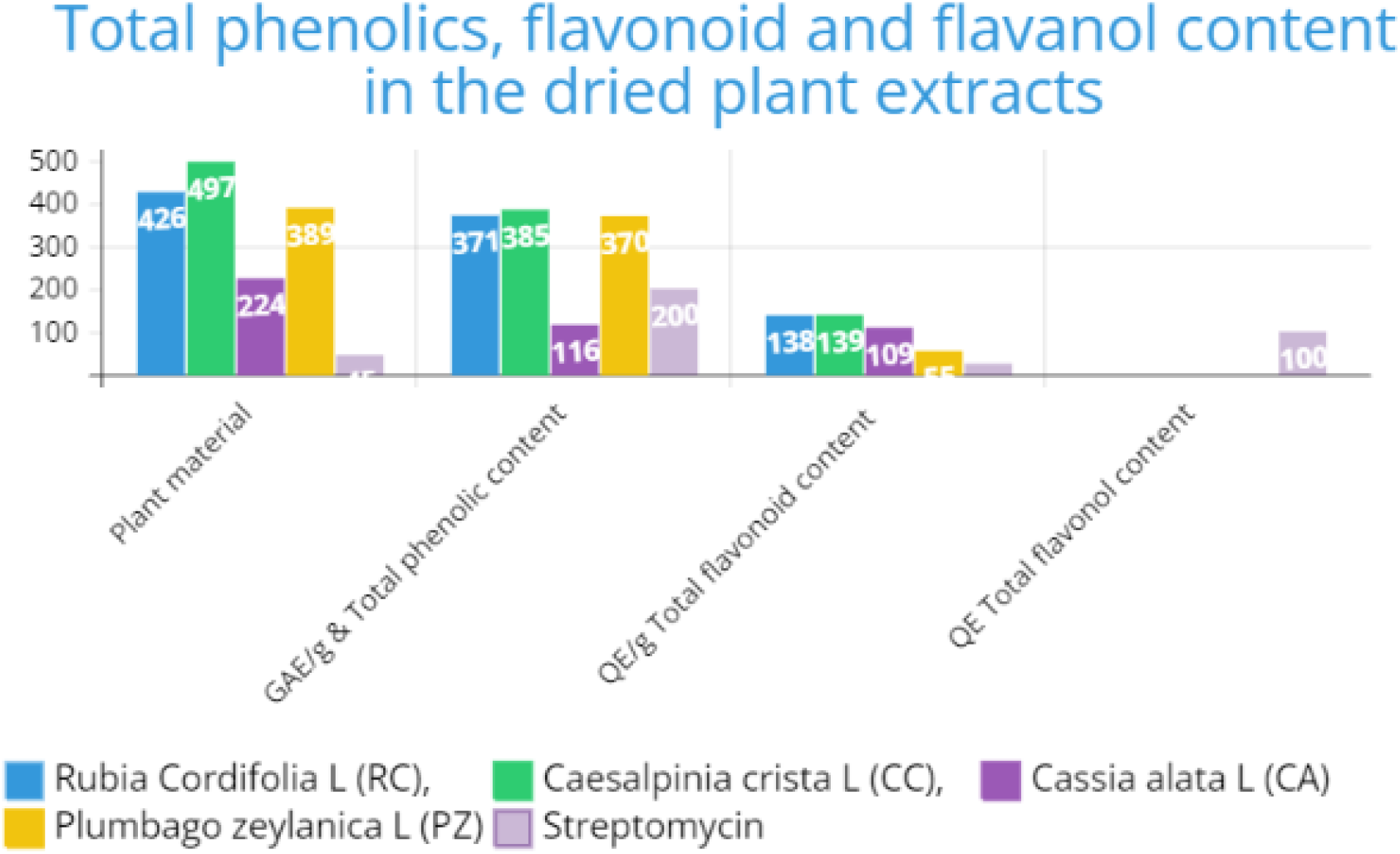

Values are means ± SD (*n* = 3). Values in the same column followed by a different letter (a-d) are significantly different (*P* < 0.05)

| Plant extracts | Zone of inhibition (mm) |  |  |  |
| --- | --- | --- | --- | --- |
|  | *A. baumannii* | *K. pneumoneae* | *P. aeruginosa* | *S. aureus* |
| Rubia Cordifolia L (RC), | 14 ± 1 | 13 ± 1.4 | 16 ± 1.2 | 15 ± 1.3 |
| Caesalpinia crista L (CC), | 18 ± 2 | 20 ± 1.5 | 23 ± 0.96 | 22 ± 1.2 |
| Cassia alata L (CA) | 14 ± 0.9 | 13 ± 1.7 | 12 ± 1.1 | 18 ± 1.3 |
| Plumbago zeylanica L (PZ) | 15 ± 1.2 | 16 ± 1.2 | 19 ± 1.3 | 17 ± 1.1 |
| Streptomycin | 17 ± 0.9 | 18 ± 0.85 | 22 ± 1.1 | 18 ± 0.86 |

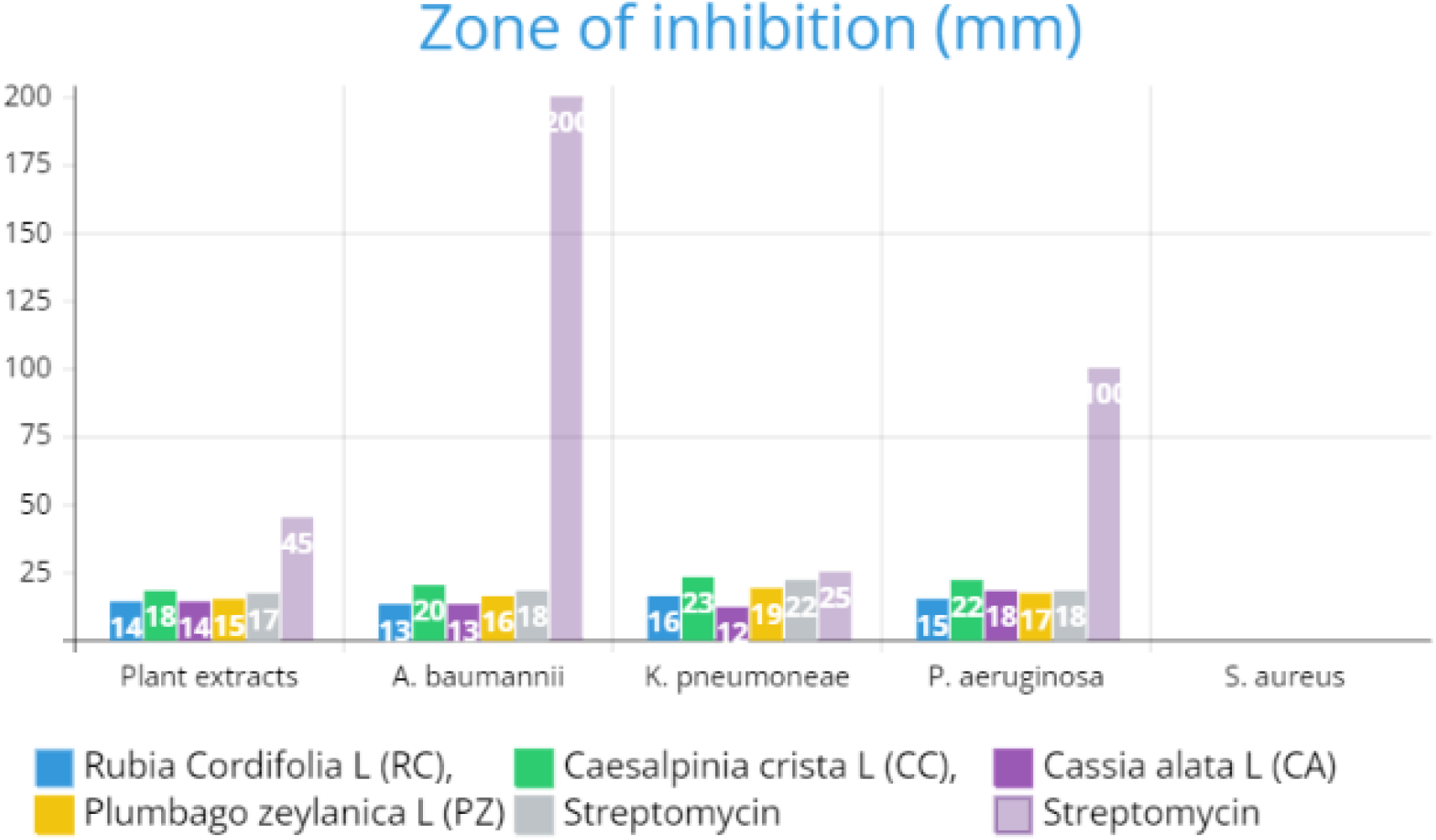

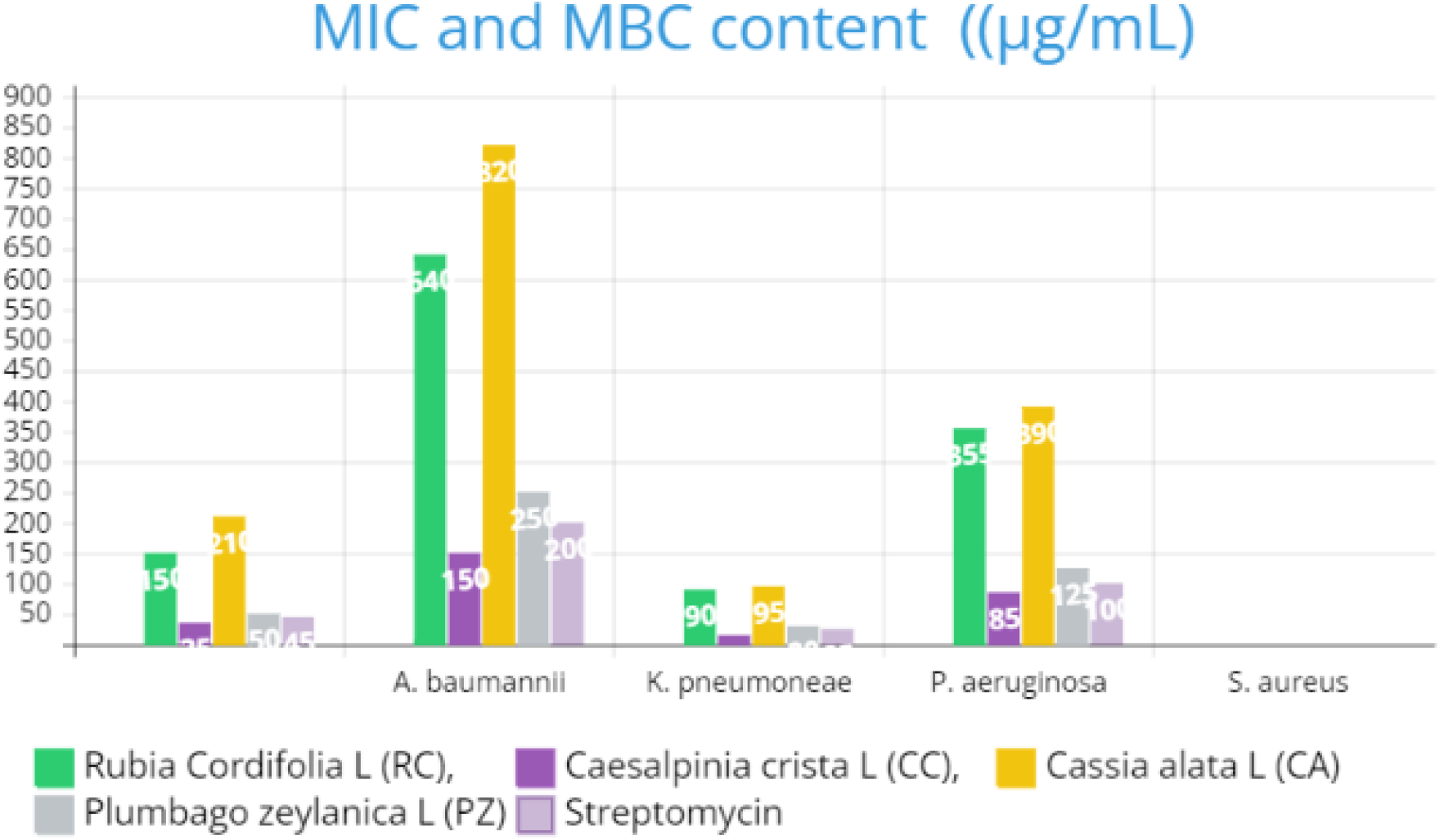

### Evaluation of Cytotoxicity

The cytotoxic potential of the methanolic extracts of the chosen plants was evaluated using brine shrimp cell cultures at three distinct concentrations: 10, 100, and 1000 mg/L. Table 6 contains the findings, which show that all of the plant extracts tested demonstrated cytotoxic action, with CA > CA > RC>CC > PZ being the ranking order of their efficiency.

**Table 6.**
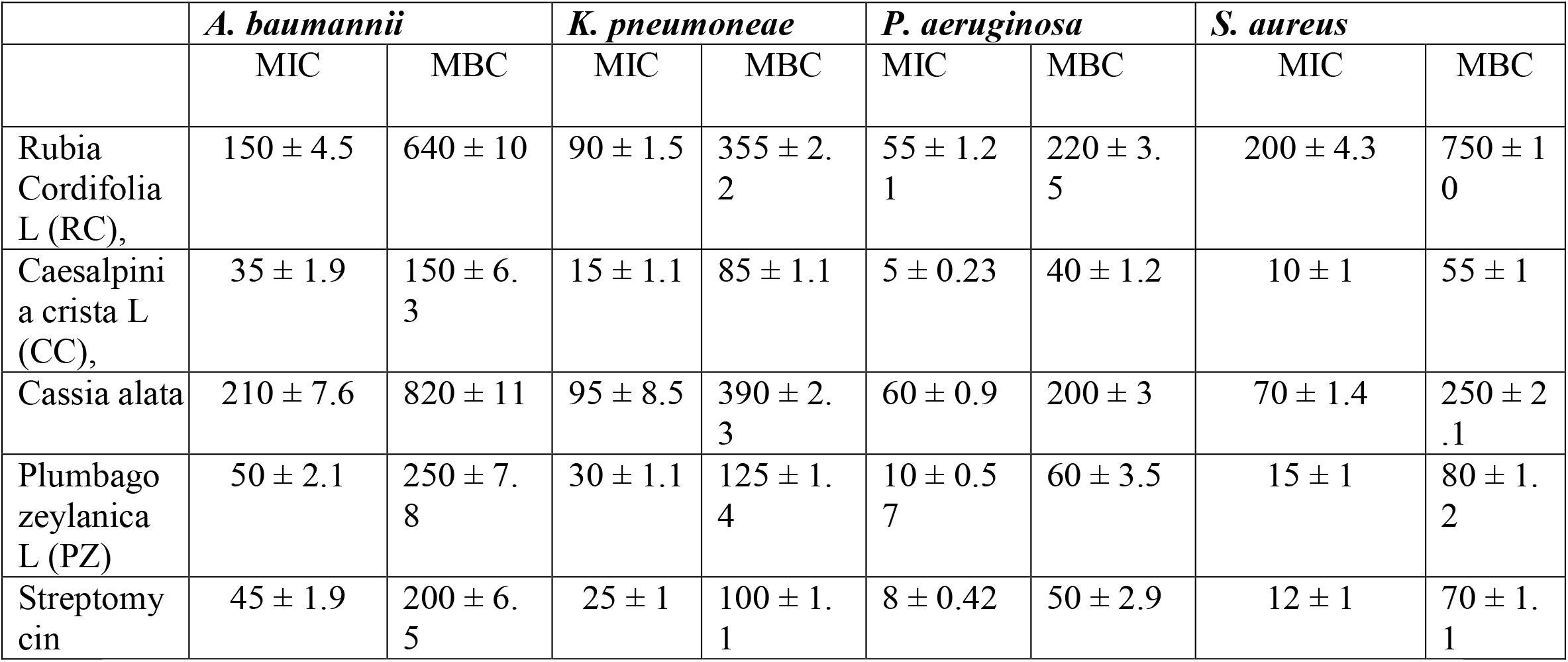
MIC and MBC content (**(μg/mL)**

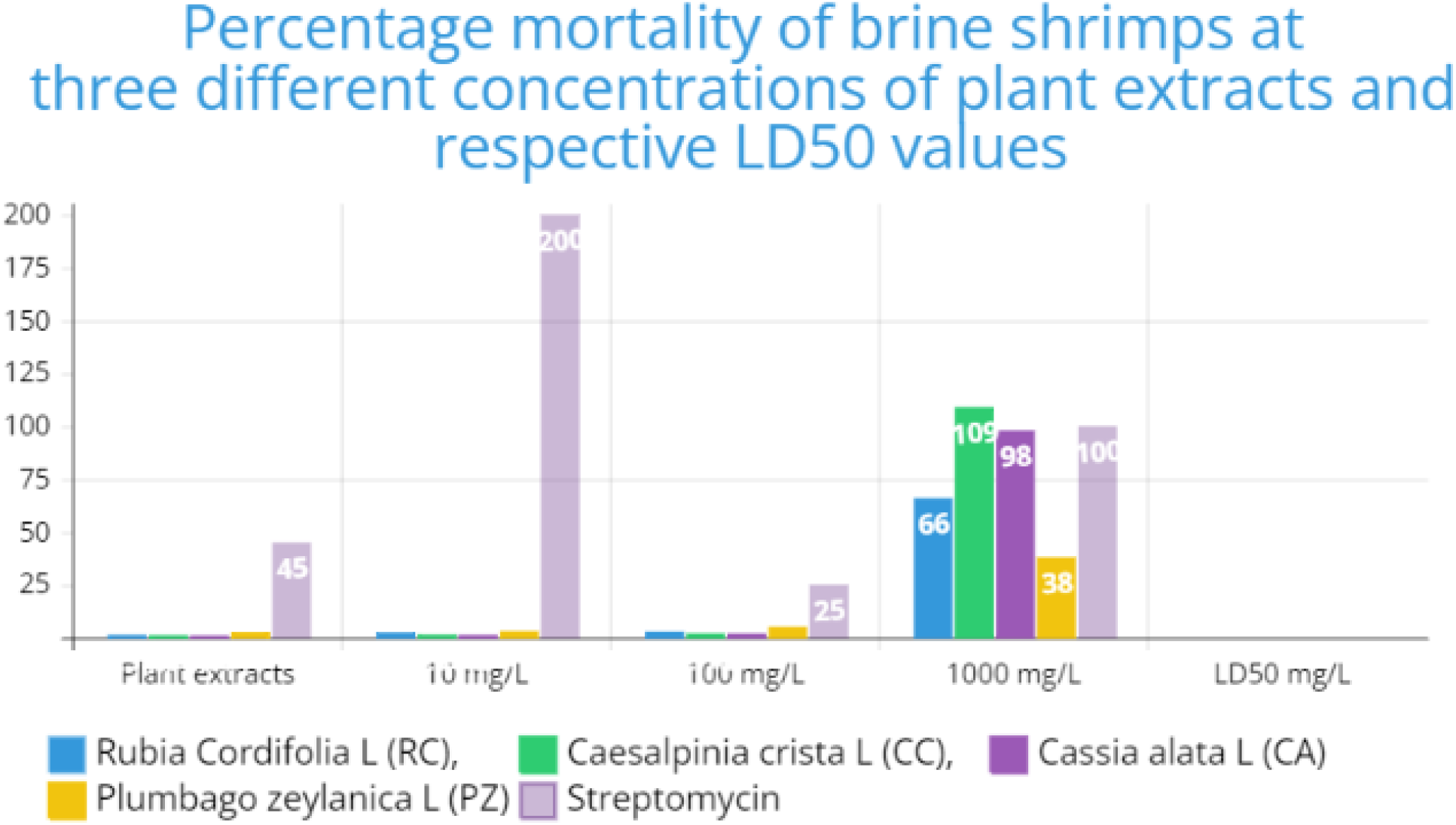

#### Anti-inflammatory and anti-proteinase activity

The suppression of albumin denaturation, haemolysis of HRBCs, and proteinase activity were measured at different doses (25–200 g/mL) of the test plant extracts in order to determine whether or not they had anti-inflammatory and anti-proteinase activity. According to the findings, proteinase activity, haemolysis of HRBCs, and albumin denaturation were all significantly suppressed by each of the extracts (Table 7). PZ was shown to prevent albumin denaturation the most effectively, followed by CA and CA, RC and CC, and then RC and CC. PZ had an IC50 value of 28 g/mL when it came to inhibiting the denaturation of albumin.

**Table 7.** Percentage mortality of brine shrimps at three different concentrations of plant extracts and respective LD_50_ values

| Plant extracts | 10 mg/L | 100 mg/L | 1000 mg/L | LD<sub>50</sub> mg/L |
| --- | --- | --- | --- | --- |
| Rubia Cordifolia L (RC), | 1.39 | 2.48 | 2.89 | 66 |
| Caesalpinia crista L (CC), | 1.23 | 1.55 | 2.13 | 109 |
| Cassia alata L (CA) | 1.09 | 1.37 | 2.09 | 98 |
| Plumbago zeylanica L (PZ) | 2.65 | 3.12 | 5.20 | 38 |

**Table 7.** Albumin denaturation, membrane protection/stabilization and proteinase inhibition potential of methanol extract of the selected plant species

| Percent inhibition |  |  |  |  |
| --- | --- | --- | --- | --- |
| Test sample | Conc. (µg/mL) | Albumin denaturation | Membrane protection | Proteinase inhibition |
|  | 25 | 29.7 ± 1.4 | 26.1 ± 0.9 | 26 ± 0.8 |
|  | 50 | 37.1 ± 1.5 | 29.4 ± 1.1 | 28.8 ± 1.1 |
| Rubia Cordifolia L (RC) | 100 | 46.8 ± 1.2 | 35.7 ± 1.0 | 35.1 ± 1.1 |
|  | 200 | 61.4 ± 1.0 | 44.6 ± 1.2 | 43.9 ± 1.5 |
| IC50 |  | 129 ± 4.6 | 247 ± 5.4 | 256 ± 5.4 |
| Aspirin |  | 84.5 ± 1.1 | 93.7 ± 1.0 | 97.1 ± 1.1 |
|  | 25 | 43.1 ± 0.8 | 31.9 ± 0.9 | 30 ± 1.0 |
| Cassia alata L (CA) | 50 | 59.5 ± 0.9 | 37 ± 1.2 | 34.8 ± 1.3 |
|  | 100 | 65.6 ± 1.1 | 47.6 ± 1.3 | 42.9 ± 1.3 |
|  | 200 | 73.3 ± 1.3 | 54.5 ± 1.3 | 52.1 ± 1.5 |
| IC50 |  | 28 ± 1.1 | 151 ± 5.0 | 175 ± 5.2 |
| Aspirin |  | 91.8 ± 1.1 | 110.9 ± 1.3 | 117.1 ± 1.3 |
|  | 25 | 17.7 ± 1.5 | 16.7 ± 1.0 | 16 ± 1.1 |
| Caesalpinia crista L (CC), | 50 | 25.3 ± 1 | 23.8 ± 1.2 | 21.9 ± 1.2 |
|  | 100 | 33.6 ± 1.5 | 24.9 ± 1.4 | 23.9 ± 1.5 |
|  | 200 | 48.7 ± 1.5 | 29.8 ± 1.4 | 27.6 ± 1.5 |
| IC50 |  | 203 ± 5.6 | 510 ± 6.6 | 577 ± 7.2 |
| Aspirin |  | 86.2 ± 0.9 | 95 ± 1.2 | 105.2 ± 1.3 |
|  | 25 | 39.1 ± 1 | 30.1 ± 1.0 | 28.5 ± 1.1 |
| Plumbago zeylanica L (PZ) | 50 | 57.6 ± 1.1 | 45 ± 1.1 | 43.6 ± 1.2 |
|  | 100 | 63.4 ± 1.3 | 59.7 ± 1.3 | 59 ± 1.3 |
|  | 200 | 71.8 ± 1.5 | 68.8 ± 1.5 | 67.9 ± 1.5 |
| IC50 |  | 43 ± 1.3 | 89 ± 2.4 | 95 ± 2.7 |
| Aspirin |  | 89.2 ± 1.1 | 95.4 ± 1.2 | 99.9 ± 1.3 |
|  | 25 | 30.7 ± 1.0 | 28.3 ± 0.8 | 27.8 ± 0.9 |

Data from Table 7 and Fig. 5 were reported as follows: Values are means ± SD (n = 3).

CA demonstrated the greatest inhibition (69%) of heat-induced hemolysis of HRBCs, followed by PZ > CA. CA was surpassed in its effectiveness by PZ. The half-maximal inhibitory concentrations (IC50) for CAF, PZ, and CAL were respectively 89, 151, and 193 g/mL. At a concentration of 200 g/mL, CAF demonstrated the highest level of anti-proteinase activity inhibition (68%), followed by PZ and CAL. The half-maximal inhibitory concentrations, or IC50 values, for CAF, PZ, and CAL were 95, 175, and 209 g/mL, respectively. As evidenced by strong correlation coefficients (R2) values of 0.76, 0.72, and 0.996 for albumin denaturation, membrane protection, and proteinase inhibition, respectively, it was discovered that there are positive connections between total phenolic content and anti-inflammatory and anti-proteinase activities. As a result, the total phenolic content of the plant extracts that were evaluated in this study may be used as an indication in order to evaluate the anti-inflammatory and anti-proteinase actions of these plant extracts.

## Discussion

The therapeutic value of the plants Rubia Cordifolia L. (RC), Cuscutapedicellata (PZ), Caesalpinia crista L. (CC), and Cassia alata L. (CA) is well-known, and all four of these species have been investigated from an ethnobotanical point of view. On the other hand, there hasn’t been enough study done on the biological activity of these plants, particularly PZ, which were gathered in the Vidisha area of Madhya Pradesh. According to the results of TLC profiling performed in a solvent system consisting of chloroform and methanol, the chosen plant extracts include a variety of chemicals. Some of these compounds include phenolic, flavonoid, tannin, terpenoid, phlobatannin, and anthraquinone. In addition to this, it was discovered that the phenolic and flavonoid content of the methanolic stem extracts of RC, PZ, CA, and CC was rather high. According to the findings of the research, the phenolic compounds of the chosen plant extracts were responsible for 93% of the antioxidant activities of those extracts. Additionally, the study discovered that favourable correlations existed between the total phenolic content of the selected plant extracts and the antibacterial and antifungal capabilities of such extracts. PZ showed the lowest MIC and MBC levels against a variety of bacterial pathogens, but the extracts of CA and CC had the highest DPPH and H2O2 radical scavenging capabilities among the three plants. The findings of this study’s antibacterial and antifungal activities may be linked to the presence of specific bioactive components such as phenolics, tannins, flavonoids, and polyphenols, respectively. These substances were investigated. The cytotoxicity and anti-inflammatory potential of the chosen plants were also evaluated as part of this research. The results revealed that the extracts of PZ, CA, and CC exhibited a considerable anti-inflammatory potential in either stabilising RBC membranes or suppressing heat-induced hemolysis. According to these findings, the chosen plant extracts contain potent anti-inflammatory, antibacterial, cytotoxic, and antioxidant effects.

In the current research, the phytochemical makeup and biological activity of four medicinal plants that are often employed in traditional medicine were analysed. The high levels of total phenolic and flavonoid content may be linked to the strong antioxidant activity that was found in all four plants, which was determined by the experiment’s findings. The antioxidant activity of the PZ extract was found to be the greatest, followed by that of the CA, CC, PZ, and RC extracts, respectively.

In addition, the anti-inflammatory, cytotoxic, and proteinase-inhibiting properties of the plant extracts were investigated as part of this research. The cells of brine prawns were shown to be susceptible to the cytotoxic effects of all four plants, with CA proving to be the most potent. In terms of their ability to reduce inflammation, every plant extract tested demonstrated reduction of albumin denaturation and hemolysis of HRBCs, with CAF showing the greatest level of proteinase activity inhibition compared to the other plant extracts.

The study of correlation revealed that there was a positive link between the total phenolic content and the observed biological activities. This conclusion was reached as a result of the findings. Based on these data, it seems that the high phenolic content of the plant extracts could be responsible for the reported biological activities of the plant extracts.

The findings of the current research provide credence to the age-old practise of making medical preparations from these plants due to the curative qualities they contain. Additional research is required to uncover the particular bioactive chemicals that are responsible for the biological activities that were observed and to evaluate whether or not these compounds have the potential to be used as natural treatments for a variety of health issues.

## Conclusion

The methanolic PZ extracts were shown to have the greatest amounts of antioxidant, antibacterial, and anti-inflammatory action, the quantities of which were substantially associated with the phenolic content of the extracts. The findings revealed that PZ stem extracts, along with CA and CA extracts, contain considerable levels of phenolic compounds. These compounds contribute to the different therapeutic actions that PZ stem extracts and CA and CA extracts are capable of, including their potential to prevent food deterioration and cure inflammation and skin irritations. In the future, research should concentrate on examining the possible application of PZ stem extracts in preventing peroxidative damage related with the development of cancer. For the purpose of determining the efficacy of these therapeutic plant extracts, notably PZ, as prospective pharmacological agents, more toxicological and phytochemical studies are required.In this research, extracts of Rubia cordifolia, Cassia alata, and Plumbago zeylanica were tested for their antibacterial, cytotoxic, and antioxidant properties. According to the findings, all three plants exhibited substantial antibacterial activity against a wide variety of harmful microorganisms, including Escherichia coli and Staphylococcus aureus. Additionally, the extracts demonstrated significant cytotoxicity against cancer cell lines, pointing to the possibility that they may be used as anticancer medicines. In addition, the extracts exhibited powerful antioxidant properties, which suggests that they may be of value in the prevention of disorders associated to oxidative stress. This research lends credence to the usage of these plants in traditional ethnomedical practises and draws attention to the plants’ potential roles as sources of medicinal chemicals.

## Notes

### Competing Interest Statement

The authors have declared no competing interest.

